# EPS8 facilitates uncoating of influenza A virus

**DOI:** 10.1101/592485

**Authors:** Gloria P. Larson, Vy Tran, Shuǐqìng Yú, Yíngyún Caì, Christina A. Higgins, Danielle M. Smith, Steven F. Baker, Sheli R. Radoshitzky, Jens H. Kuhn, Andrew Mehle

## Abstract

All viruses balance interactions between cellular machinery co-opted to support replication and host factors deployed to halt the infection. We used gene correlation analysis to perform an unbiased screen for host factors involved in influenza A virus (FLUAV) infection. Our screen identified the cellular factor epidermal growth factor receptor pathway substrate 8 (EPS8) as the highest confidence pro-viral candidate. Knockout and overexpression of EPS8 confirmed its importance in enhancing FLUAV infection and titers. Loss of EPS8 did not affect virion attachment, uptake, or fusion. Rather, our data show that EPS8 specifically functions during virion uncoating. EPS8 physically associated with incoming virion components, and subsequent nuclear import of released ribonucleoprotein complexes was significantly delayed in the absence of EPS8. Our study identified EPS8 as a host factor important for uncoating, a crucial step of FLUAV infection during which the interface between the virus and host is still being discovered.

## INTRODUCTION

Attachment and entry into a host cell is the first bottleneck virions encounter during infection. Virion entry requires efficient use of the host cell environment while simultaneously evading cellular immune responses. Influenza A virus (FLUAV; *Orthomyxoviridae*: *Alphainfluenzavirus*), like all viruses, largely depends upon existing cellular machinery to successfully complete these initial stages of infection.

During the first step of infection, attachment, FLUAV hemagglutinin (HA) binds to the target cell via sialic acid linkages on host glycoproteins (Dou et al., 2018). Virions are internalized via receptor-mediated endocytosis and less frequently through an alternative macropinocytosis pathway (Matlin et al., 1981; de Vries et al., 2011). Once within endosomes, virions are trafficked towards the nucleus using the cytoskeletal components actin, dynein, and microtubules (Lakadamyali et al., 2003). The endosome matures and acidifies during cellular trafficking, and the virion interior is also acidified through the function of the viral ion channel M2 (Pinto et al., 1992). The low pH in the endosome causes conformational changes in HA that drive fusion of the viral and endosomal lipid membranes, while low pH within the virion causes the viral matrix protein M1 to dissociate from the inner membrane of the viral envelope (Bukrinskaya et al., 1982; Maeda and Ohnishi, 1980; Martin and Helenius, 1991; Zhirnov, 1990). Fusion of the two membranes releases a capsid-like viral core consisting of viral ribonucleoproteins (vRNPs) enclosed in an M1 shell-like structure into the cytoplasm. This complex engages the cellular aggresome to complete uncoating, and the released vRNPs are imported into the nucleus by cellular karyopherins (Banerjee et al., 2014; Melen et al., 2003; O’Neill et al., 1995; Wang et al., 1997). Once in the nucleus, a pioneering round of transcription occurs on the incoming vRNPs that initiates replication and secondary rounds of transcription of the viral genome.

High-throughput screening approaches have expanded our knowledge of specific cellular cofactors involved in FLUAV infection, with many of these methods identifying host factors involved in viral entry. Gene disruption screens identified host factors involved in sialic acid metabolism utilized for attachment (Carette et al., 2009; Han et al., 2017). Vacuolar ATPases involved in endosomal acidification and other host factors facilitating fusion and uncoating were identified through siRNA knockdown, proteomic, and overexpression screens (Banerjee et al., 2014; König et al., 2010; Lee et al., 2017; Mar et al., 2018; Yángüez et al., 2018). These studies also revealed previously unknown steps of FLUAV particle entry such as the role of the aggresome in viral uncoating (Banerjee et al., 2014). Frequent identification of cellular factors involved in viral entry highlights the critical role of this process during viral replication. Despite these discoveries, however, the mechanistic details of steps occurring after fusion remain poorly understood.

Here, we conducted a screen using gene correlation analysis to identify host factors involved in FLUAV infection. Gene correlation analysis exploits naturally occurring variations in gene expression across multiple cell lines without the need to exogenously manipulate the cellular environment. Variations in cellular gene expression were used to identify factors affecting a phenotype of interest, in this case susceptibility to FLUAV infection. We identified epidermal growth factor receptor (EGFR) pathway substrate 8 (EPS8) as a pro-viral cellular cofactor during the early stages of infection. We confirmed that EPS8 enhances FLUAV gene expression and replication, whereas knockout of EPS8 reduced susceptibility to infection. Step-wise dissection of the viral entry process revealed that EPS8 specifically facilitates uncoating of the viral core. Thus, we identified EPS8 as an important component of the FLUAV uncoating process, a necessary step for successful viral genome transcription and replication.

## RESULTS

### Gene correlation analysis identifies putative enhancers and suppressors of FLUAV replication

To overcome limitations of previous screening methodologies, we sought to identify both enhancers and suppressors of FLUAV replication in an unbiased manner. We utilized gene correlation analysis which relied on inherent differences in gene expression among different cell lines and consequently did not require external manipulation of the cellular environment. The National Cancer Institute-60 (NCI-60) panel consists of 59 distinct cell lines with well-characterized transcriptomic profiles (Shankavaram et al., 2007; Weinstein and Pommier, 2003). The diversity of cell types and the depth of transcriptomic data permit high confidence genome-wide correlations between cellular gene expression and infection susceptibility (Kondratowicz et al., 2013; Lenaerts et al., 2012; Schowalter et al., 2012). We therefore inoculated the NCI-60 panel of cell lines with a single-cycle variant of A/WSN/1933 (H1N1, WSN) encoding GFP (WSN-GFP) (Figure 1A). Using WSN-GFP, we specifically focused on host factors involved in early stages of infection up to and including viral gene expression and translation.

**Figure 1.**
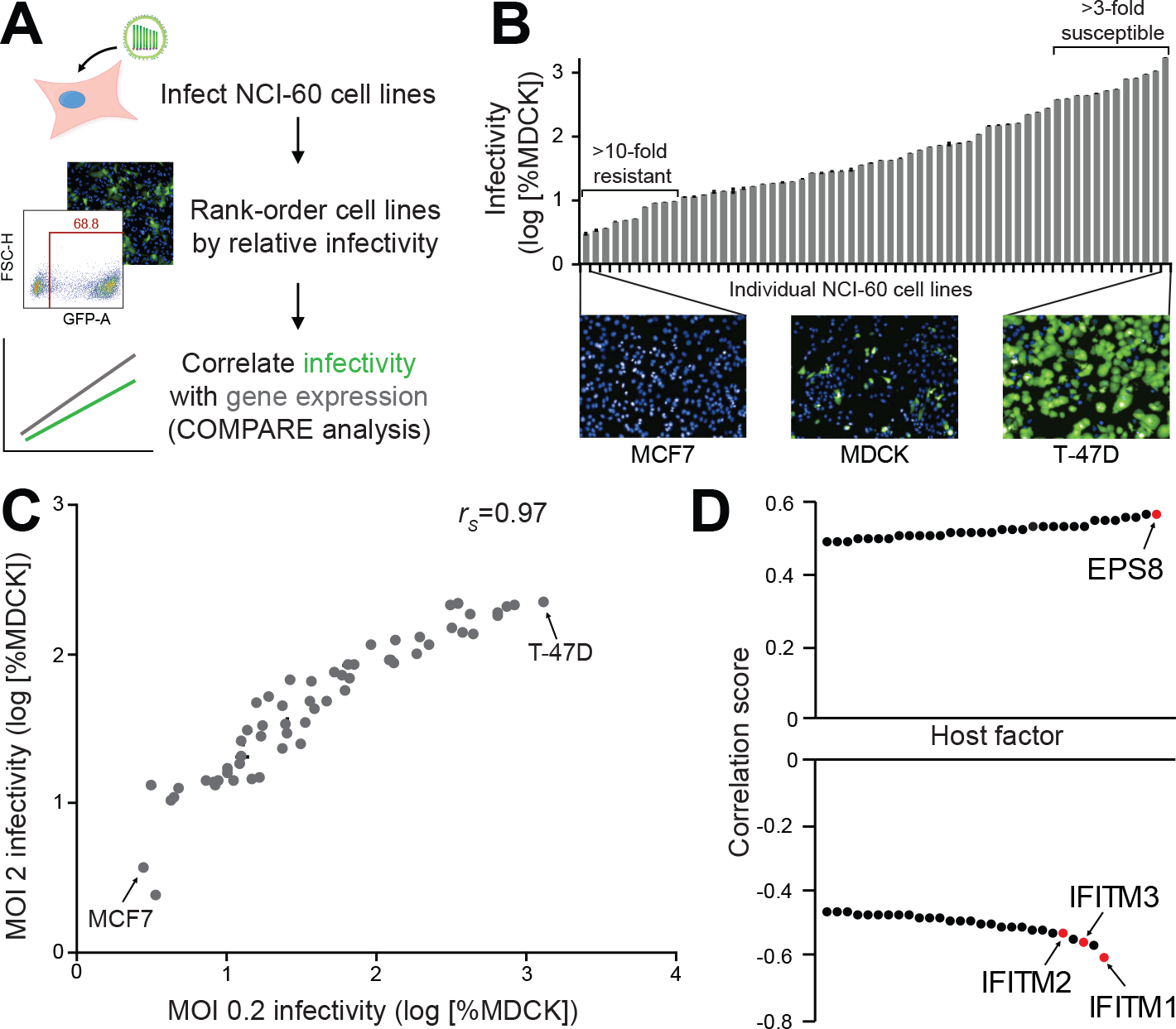
Gene correlation analysis identifies putative enhancers and suppressors of FLUAV replication. (A) Experimental workflow for NCI-60 screen. NCI-60 cell lines were inoculated with FLUAV encoding GFP, infections were visualized by fluorescence microscopy and quantified by flow cytometry at 24 hpi, and data were normalized to control MDCK cells inoculated in parallel. (B) Infectivity at an MOI of 0.2 was determined relative to MDCK cells (mean of n=3 ± SD). Images of highly resistant (MCF7) and hypersensitive (T-47D) infected cell lines are shown compared to the control MDCK cells. (C) Pairwise comparison of replicate NCI-60 screens performed at an MOI of 0.2 or 2. (mean of n=3 ± SD, rs = Spearman’s correlation co-efficient) (D) COMPARE analysis of MOI 0.2 infectivity data identified top hits for putative pro-viral and anti-viral factors. See also Figure S1.

The permissiveness of each cell line to WSN-GFP was determined and rank-ordered relative to Madin-Darby canine kidney (MDCK) cells, which are frequently used for the propagation of FLUAV (Figures 1A and 1B; Table S1). We detected a broad range of susceptibility when infecting cells at a multiplicity of infection (MOI) of 0.2. Relative to MDCK cells, 10 cell lines were highly refractory to WSN-GFP (at least a 10-fold decrease in infection rate) and 12 cell lines were highly permissive (at least a 3-fold increase in infection rate). There was no obvious association between susceptibility, cell type, tumor type, or tissue of origin. MCF7 breast tumor cells were the most refractory with a normalized infection rate of only about 3%, whereas T-47D cells, another breast tumor cell line, were the most susceptible with an infection rate of approximately 1300%. These data were highly reproducible with a strong correlation between results from two independent replicate screens (Figure S1A). To ensure that the assay captured the full dynamic range of susceptibility, especially for the highly resistant cell lines, the screen was repeated at an MOI of 2 (Figure S1B). Similar infectivity trends were detected at both MOIs, although the upper limit of the assay was reached for multiple cell lines at the higher MOI where effectively all cells were infected (Figure 1C and S1C). The number of infected cells increased at the higher MOI for most of the resistant cell lines, indicating that these cell lines are not completely refractory to FLUAV infection (Figure S1D).

The broad distribution of infectivity across the NCI-60 panel suggested that cell-intrinsic differences impacted susceptibility to FLUAV infection. To identify cellular factors impacting FLUAV susceptibility, we calculated linear pairwise correlation coefficients between host gene expression within the NCI-60 panel of cell lines and susceptibility to infection using the COMPARE algorithm (Zaharevitz et al., 2002). We identified top hits for putative enhancers or suppressors of FLUAV infection based on their strong correlation scores (Figure 1D; Table S2). Host genes identified as putative enhancing factors exhibited expression patterns that paralleled susceptibility to infection, yielding a positive correlation score. Conversely, expression of host genes identified as putative suppressive factors was inversely related to susceptibility, resulting in a negative correlation score. Notably, some of our strongest hits for suppressors of FLUAV infection were the interferon-inducible transmembrane proteins (IFITMs). IFITM1, IFITM2, and IFITM3 have previously been characterized as potent inhibitors of FLUAV infection, providing confidence in our approach (Figure 1D) (Brass et al., 2009). Most other candidate genes, including EPS8, have not been previously associated with FLUAV susceptibility, revealing that gene correlation analysis can identify new host factors that regulate FLUAV infection.

### EPS8 enhances FLUAV gene expression and titers

The putative enhancer with the strongest correlation score was EPS8, an adaptor protein involved in signaling via the epidermal growth factor receptor (EGFR) and other pathways as well as modulating of actin dynamics (Figure 1D) (Di Fiore and Scita, 2002; Hertzog et al., 2010). To validate the results of the screen and confirm a pro-viral function for EPS8, we assessed the effect of EPS8 on viral gene expression and replication. EPS8 was transiently overexpressed in human embryonic kidney 293T cells and infected with a replication-competent reporter version of WSN (WSN PASTN) to quantitatively measure viral gene expression (Tran et al., 2013). EPS8 overexpression increased viral gene expression during infection nearly two-fold relative to the empty vector control (Figure 2A). Endogenous and overexpressed EPS8 levels were confirmed by immunoblot. We then assayed viral titers when EPS8 was stably overexpressed in human lung epithelial A549 cells. Viral titers 24 hours post-infection (hpi) were increased by over 15 fold in stable EPS8-overexpressing cells relative to wild type (WT) cells (Figure 2B). Thus, overexpression of EPS8 enhances infection and replication in two different human cell lines, confirming the pro-viral correlation identified in the screen.

**Figure 2.**
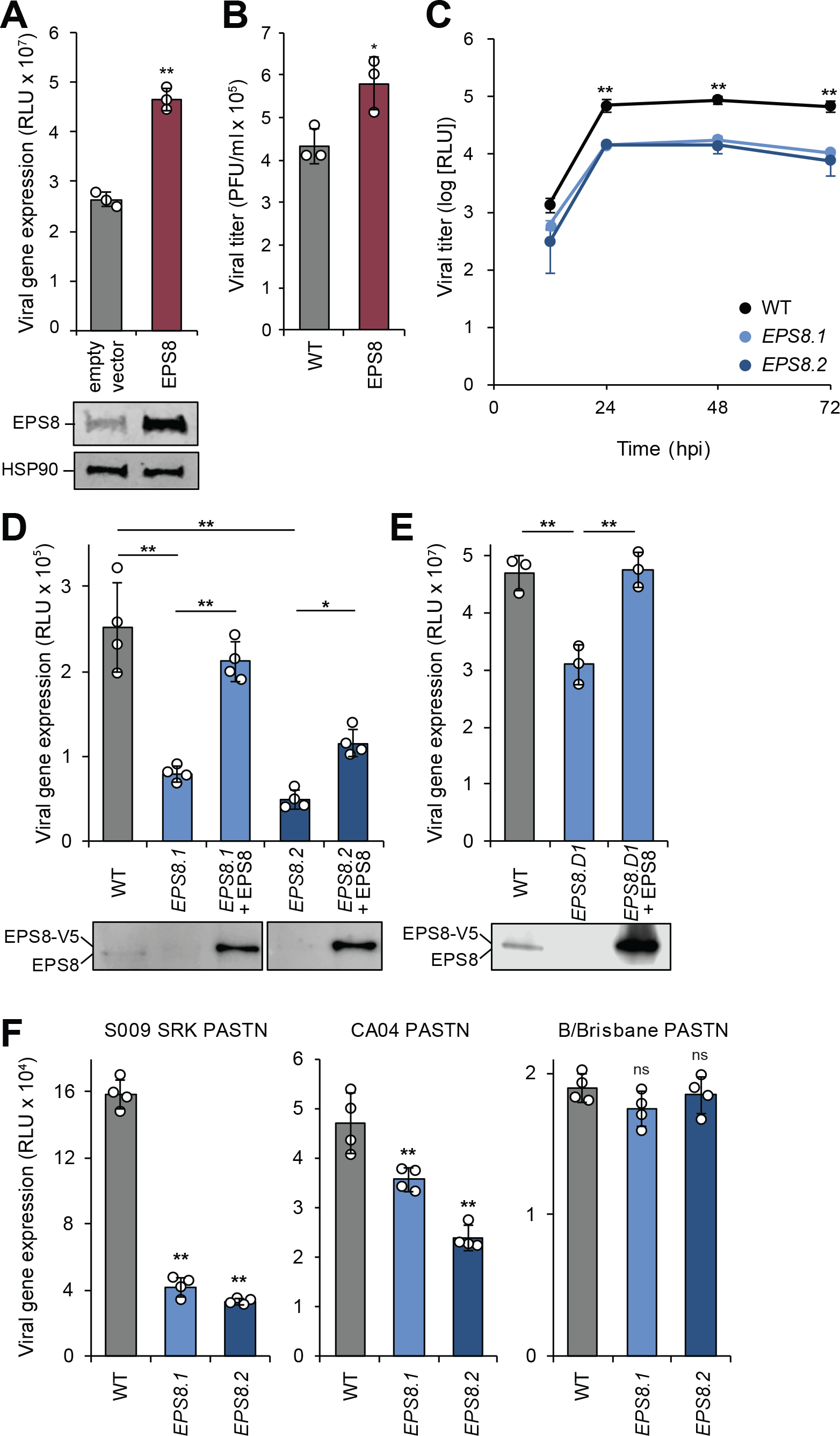
EPS8 enhances FLUAV gene expression and titers. (A) 293T cells transiently overexpressing EPS8 were infected with WSN PASTN and viral gene expression assayed (mean of n=4 ± SD). EPS8 expression was confirmed by immunoblot. (B) A549 cells stably overexpressing EPS8 were infected with WSN and viral titer assayed at 24 hpi (mean of n=3 ± SD). (C) EPS8-edited A549 cells were infected with WSN PASTN, virus was harvested at the indicated times, and titers were assessed using luciferase activity (mean of n=3 ± SD). (D-E) EPS8-edited and complemented A549 (D) or 293 (E) cells were infected with WSN PASTN to assay viral gene expression (mean of n=4 ± SD). EPS8 expression was confirmed by immunoblot. (F) Viral gene expression was assayed in EPS8-edited cells infected with S009 SRK PASTN, CA04 PASTN, or B/Brisbane PASTN (mean of n=4 ± SD). * = p<0.05, ** = p<0.01, ns = not significant. (A-B) were analyzed by Student’s two-tailed t-test, unequal variance. Multiple comparisons were made in (C-F) using a one-way ANOVA with post-hoc Tukey HSD test when compared to wild type A549 cells. See also Figure S2, S3, and S4.

We next used CRISPR-Cas9 to generate clonal EPS8 knockout A549 cells. Sanger sequencing confirmed genotypic changes predicted to result in knockout of EPS8 in two independent clonal lines (*EPS8.1*, *EPS8.2*) (Figure S2A and S2B). Immunoblotting for endogenous EPS8 revealed a dramatic reduction in EPS8 protein levels but not a complete loss in our edited clones (Figure S2C). *EPS8.1* retained about 25% of the amount of EPS8 observed in the parental cells, whereas *EPS8.2* levels were nearly undetectable. Editing occurred adjacent to the splice donor in exon 2 of *EPS8*, raising the possibility that alternative splice donors may be exploited to support the low levels of EPS8 protein expression detected (Figure S2B and S2D). These cell clones were used to further examine the importance of EPS8 during FLUAV infection.

Viral replication and gene expression were assayed in the *EPS8*-edited cells. Both *EPS8.1* and *EPS8.2* cell lines had defects in multicycle replication and viral gene expression assays. Viral titers were reduced by about 10-fold in both *EPS8*-edited lines compared to parental cells (Figure 2C). Viral gene expression was reduced 4-5 fold in A549 cells with edited *EPS8* relative to wild type cells (Figure 2D). The decrease in viral gene expression was more pronounced in *EPS8.2*, the cell line with the lower level of EPS8 expressed. Stable complementation with EPS8 rescued viral gene expression in both edited lines (Figure 2D), suggesting the defects in gene expression were specifically due to decreases in EPS8 levels. To obtain a true knockout phenotype, *EPS8* was edited in 293 cells (Supplemental Figure 3). EPS8 knockout 293 cells (*EPS8.D1*) exhibited a significant decrease in viral gene expression, which was restored by transient complementation (Figure 2E). *EPS8* editing or knockout thus decreases viral gene expression in two different cell lines.

**Figure 3.**
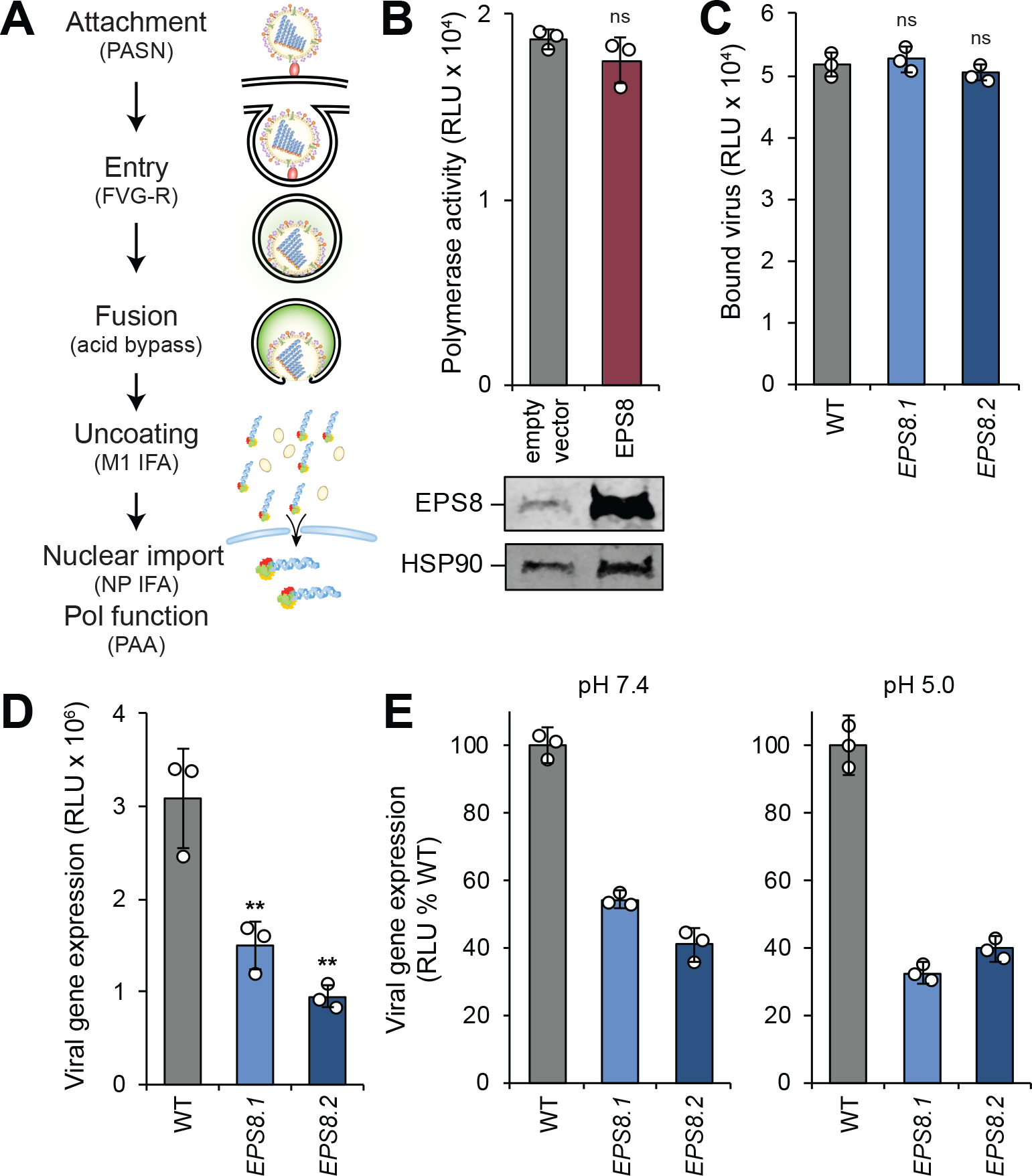
EPS8 functions post-fusion but before viral gene expression during FLUAV infection. (A) Early stages in the FLUAV replication cycle were systematically probed with the indicated reagents or assays detailed in the text. PASN = bioluminescent virions. FVG-R = recombinant FLUAV expressing VSIV-G. IFA = immunofluorescence assay. PAA = polymerase activity assay. (B) Polymerase activity assays were performed in 293T cells expressing RNP components with or without exogenous EPS8. EPS8 expression was confirmed by immunoblot. (C) Virion attachment was assayed in EPS8-edited cells incubated with bioluminescent PASN. (D) Wild type or EPS8-edited cells were inoculated with FVG-R and viral gene expression was measured 8 hpi. (E) Acid bypass assays were performed on virions attached to wild type of EPS8-edited cells. Cells were transiently treated with buffers at physiological pH 7.4 to initiate canonical viral entry, or acidic pH 5.0 to cause fusion at the cell surface. Viral gene expression was measured 8 h after treatment. For all, data are mean of n=3 ± SD. ** = p<0.01, ns = not significant. (B) was analyzed by Student’s two-tailed t-test, unequal variance. Multiple comparisons were made in (C-E) using a one-way ANOVA with post-hoc Tukey HSD test when compared to wild type A549 cells.

Given that both cell types exhibited similar phenotypes, we continued our investigation using the edited A549 cell lines, as these cells are of lung origin and more closely represent natural target cells during influenza virus infection. We assessed whether EPS8 affected primary influenza virus isolate infection. Cells were inoculated with a reporter virus encoding an avian-background RNP A/green-winged teal/OH/175/1983 in a WSN backbone (S009 SRK PASTN; H2N1) or a reporter version of the primary isolate A/California/04/2009 (CA04 PASTN; H1N1). We also infected *EPS8*-edited cells with the influenza B virus (FLUBV) primary isolate B/Brisbane/60/2008 (B/Brisbane PASTN) (Figure 2F). Consistent with the results obtained using WSN, editing of *EPS8* reduced viral gene expression for the influenza A strains S009 SRK PASTN and CA04 PASTN. Interestingly, *EPS8* editing did not affect B/Brisbane PASTN gene expression (Figure 2F). We explored infection specificity further by assessing the relative infection rates of A549 cells overexpressing EPS8 in response to challenge by diverse viruses (Figure S4). EPS8 expression levels did not alter infection rates of Marburg virus (MARV) or Junín virus (JUNV). In contrast, EPS8 overexpression caused decreased Ebola virus (EBOV), Venezuelan equine encephalitis virus (VEEV), and Rift Valley fever virus (RVFV) infection rates. Hence, altering EPS8 expression does not generically affect viral replication. Together, these data confirm that EPS8 acts as a pro-viral host factor during FLUAV infection and exhibits specific effects on cell infectivity depending on the virus.

### EPS8 functions post-fusion but before viral gene expression during FLUAV infection

The structure of the NCI-60 screen and our data indicated EPS8 functions in early stages of FLUAV replication. We therefore conducted a series of experiments to determine where in the viral replication cycle EPS8 functioned to enhance infection (Figure 3A). We first assessed whether EPS8 affects infection through a mechanism that directly impacts viral polymerase activity. Polymerase activity was reconstituted in the absence of infection by expressing the heterotrimeric viral polymerase subunits PA, PB1, and PB2, nucleoprotein (NP), and a vRNA-like reporter encoding firefly luciferase. Polymerase activity was not statistically different in the presence or absence of exogenous EPS8 (Figure 3B). Immunoblotting confirmed high levels of exogenously expressed EPS8. This finding establishes that EPS8-mediated enhancement of viral gene expression is not due to direct impacts on the viral polymerase but rather an upstream step in the early stages of infection.

We probed each successive step that occurs early in the infectious cycle, beginning with viral attachment. Wild-type or edited cells were incubated with bioluminescent virions (PASN) that package nanoluciferase into the viral particle (Tran et al., 2015). Cells were incubated at 4°C to enable binding but prevent internalization of virions, and luciferase activity was assayed from the bound virions. There was no statistical difference in the amount of virus bound to wild type and both *EPS8*-edited cell lines, indicating EPS8 is not necessary for FLUAV attachment to cells (Figure 3C). To ascertain if EPS8 affects HA-mediated entry or the fusion process, we infected cells with FLUAV encoding a different entry protein, FVG-R, a recombinant virus expressing vesicular stomatitis Indiana virus glycoprotein (VSIV-G) instead of HA (Hao et al., 2008). Viral gene expression decreased in *EPS8*-edited cells infected with FVG-R compared to wild type cells (Figure 3D). This observed decrease in viral gene expression was similar to the decrease demonstrated during infection with *bona fide* FLUAV (Figure 2D and 2E) and suggests EPS8 does not specifically target HA-mediated entry.

Following attachment and entry, FLUAV traffics in an endosome that undergoes acidification which results in fusion of the endosomal and viral membranes. The function of EPS8 during endosomal acidification and fusion was tested using an acid bypass assay. Acid bypass replaces the canonical entry route with fusion of viral and plasma membranes at the cell surface, depositing vRNPs into the cytoplasm where subsequent steps of infection then proceed as usual (Banerjee et al., 2014; Matlin et al., 1981). As in the attachment assay, virions bound to the surface of wild type or edited cells at 4°C to synchronize infection. Cells were shifted to 37°C and transiently held at acidic conditions (pH 5.0) to initiate fusion at the cell surface or held at physiological conditions (pH 7.4) permitting canonical entry to proceed as a control. *EPS8* editing resulted in a decrease in viral gene expression when infections were initiated at physiological pH (Figure 3E), consistent with prior data showing defects in gene expression during unsynchronized infections (Figure 2D and E). Bypassing canonical entry by treating cells with acidic conditions did not restore viral gene expression in the edited cells (Figure 3E), indicating EPS8 does not function during endosomal acidification. Together, these data establish that the effects of EPS8 during FLUAV infection are independent of virion attachment, endosomal entry, and HA-mediated fusion.

### EPS8 is crucial for viral uncoating

Our line of experimentation indicated EPS8 functions at a step following release of the viral core into the cytoplasm but before viral gene expression. Therefore, we considered whether EPS8 facilitates viral uncoating. This process can be quantified by visualizing the redistribution of punctate matrix protein (M1) staining of intact particles to diffuse staining of M1 released throughout the cytosol (Figure 4A and S5A) (Banerjee et al., 2013). Wild type and *EPS8*-edited cells were synchronously infected, and M1 localization was quantified at various times post-inoculation. As expected, most M1 staining was punctate in wild type cells early in infection and then became diffuse at 1.5 hpi (Figure 4B). By contrast, uncoating was greatly delayed in both cell lines where *EPS8* was edited. Diffuse M1 staining was detected in only 10-15% of *EPS8*-edited cells at 1.5 hpi compared to successful uncoating in almost all wild type cells at the same time point. Following release from the endosome, viral cores are trafficked to the aggresome to complete uncoating (Banerjee et al., 2014). Co-precipitations were used to probe how EPS8 might function during this period. Synchronized infections were initiated on *EPS8*-edited cells stably complemented with wild type EPS8. NP specifically co-precipitated with EPS8, suggesting EPS8 physically interacts with incoming viral cores (Figure 4C). These data implicate EPS8 as an important host factor during viral uncoating.

**Figure 4.**
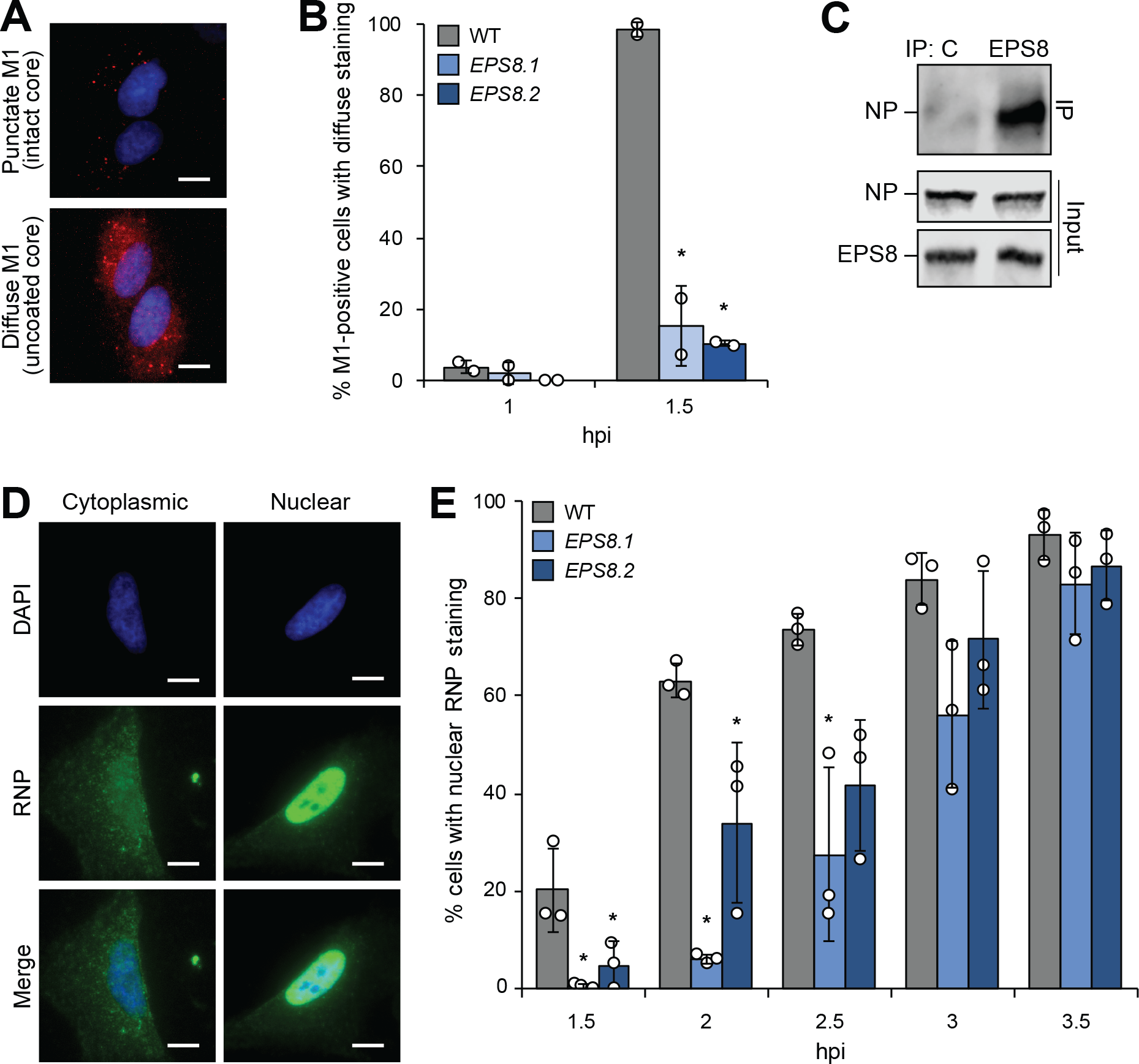
EPS8 is crucial for viral uncoating. (A) A549 cells synchronously infected with WSN were stained for M1 (red) and the nucleus (blue). Representative images show punctate M1 consistent with intact viral cores and diffuse M1 staining that occurs following viral uncoating. (B) Quantification of diffuse staining in M1-positive cells (mean of n=2 ± SD). (C) EPS8-edited A549 cells complemented with EPS8 were infected and lysates subjected to immunoprecipitation. Co-precipitating NP and total NP and EPS8 expression were confirmed by immunoblot. (D) Wild type A549 cells infected with WSN were stained for viral RNPs (green). Representative images show cytoplasmic RNP staining or nuclear RNP staining determined by colocalization with the nucleus (blue). (E) Quantification of the number of cells with nuclear RNP staining at each time point (mean of n=3 ± SD). * = p<0.05 one-way ANOVA with post-hoc Tukey HSD test when compared to wild type A549 cells. Scale bar = 20 μm. See also Figure S5.

Uncoating releases vRNPs into the cytosol where they are subsequently imported into the nucleus prior to viral gene expression. The defects in uncoating we detected in *EPS8*-edited cells predict that these cells should also exhibit delayed nuclear import. To test this possibility, we again used synchronized infections and immunofluorescence to examine the subcellular localization and kinetics of vRNP nuclear import over time. Staining for NP, the major protein component of vRNPs, revealed characteristic cytoplasmic localization of incoming vRNPs early during infection followed by distinct nuclear localization (Figure 4D). Discrete cytoplasmic and nuclear localizations of vRNPs were also detected in *EPS8*-edited cells (Figure S5B). Nuclear-localized vRNPs were detected in wild type cells as early as 1.5 hpi and the number of cells with nuclear vRNP staining increased over time, consistent with the timing of viral uncoating reported above (Figure 4E). Cells lacking wild type levels of EPS8, however, exhibited significantly delayed kinetics of nuclear import. Compared to wild type cells, import rates in *EPS8*-edited cells were delayed by 1 hour. This trend continued until 3.5 hpi when import in edited cells finally matched that of wild type cells (Figure 4D). While import was delayed in edited cells, it followed a similar trajectory to wild type cells once initiated, suggesting that vRNP import was not directly altered by changes to EPS8 expression. Thus, defects in uncoating (Figure 4B) result in delayed nuclear import (Figure 4E) and ultimately a reduction in viral gene expression (Figure 2D), reinforcing the conclusion that EPS8 is a key component of the cellular machinery utilized for viral uncoating.

## DISCUSSION

Through gene correlation analysis, we conducted an unbiased genome-wide screen to identify host factors that have a functional impact on early stages of FLUAV replication. Our highest confidence pro-viral candidate was EPS8, a cytoplasmic protein involved in EGFR signaling and regulation of actin dynamics. We showed that EPS8 expression enhanced viral gene expression and titers, whereas loss of wild-type EPS8 caused defects in gene expression and viral replication. Step-wise investigation of the early stages of infection revealed that EPS8 functions independent of virion attachment, endosomal acidification, or HA-dependent fusion. Rather, EPS8 specifically functioned during the uncoating of the incoming viral cores. Defects in viral uncoating slowed the kinetics of vRNP nuclear import in *EPS8*-edited cells, corresponding with the overall delay in viral gene expression and replication in these cells. These data support a new role for EPS8 as a cofactor important for viral uncoating during FLUAV infection.

The host factors used during FLUAV uncoating are not yet fully understood. Uncoating begins in the maturing endosome where the drop in pH opens the M2 ion channel in the viral membrane (Pinto et al., 1992). The influx of potassium ions and protons into the virion interior initiates conformational changes that relax interactions between the matrix protein M1 and vRNPs, making the core competent for uncoating and disassembly of the RNP bundle (Stauffer et al., 2014). Following fusion of the viral and host membranes, the core requires further processing to fully disassemble. Unanchored ubiquitin chains packaged within the virion help direct the core to the cellular aggresome where mechanical forces have been proposed to accelerate uncoating and release of vRNPs into the cytosol (Banerjee et al., 2014). Our data now implicate EPS8 as another host factor important during these later stages of uncoating.

EGFR has previously been implicated in FLUAV entry during virion internalization (Eierhoff et al., 2010). Our data, however, indicate that loss of wildtype EPS8 does not alter attachment and that bypassing internalization by forcing fusion at the plasma membrane does not rescue defects in these cells (Figure 3), suggesting EPS8 activity is independent of EGFR signaling. EPS8 is also involved in modulating actin dynamics (Hertzog et al., 2010). Actin has been implicated in the rapid movement of virion-containing endosomes immediately after virion internalization and also plays a role in the discrete steps post-fusion but before uncoating is completed (Banerjee et al., 2014; Lakadamyali et al., 2003). A role for actin during post-fusion uncoating is the same step where our data revealed EPS8 functions, raising the possibility that it is the ability of EPS8 to engage and modulate actin dynamics that is important for uncoating.

Although cells lacking EPS8 have decreased FLUAV gene expression, that was not the case during FLUBV infection. FLUAV and FLUBV are structurally similar, and it is tempting to generalize that the replication cycle is largely the same for the two viruses. Like FLUAV, FLUBV utilizes receptor-mediated endocytosis for entry (Shaw and Palese, 2013). Acidification of the FLUBV virion interior is facilitated by viral membrane protein and proton pump BM2, a FLUAV M2 homolog (Mould et al., 2003). FLUBV undergoes uncoating after fusion of viral and endosomal membranes, but many of the details of FLUBV uptake and uncoating are still unknown. Interestingly, cellular immune responses to FLUBV infection differ from those to FLUAV infection (Jiang et al., 2016; Mäkelä et al., 2015). Therefore, it is possible that host processes involved in other steps of FLUBV infection also differ, as suggested by the discordant importance of EPS8 for FLUAV and FLUBV.

We also considered the possibility that other viruses using receptor-mediated endocytosis or similar internalization pathways could be affected by EPS8. A panel of RNA viruses using diverse cellular receptors and entry mechanisms was used to infect A549 cells overexpressing EPS8 (Figure S4). There were no obvious trends or associations with viral families or entry pathways. Nonetheless, these results indicate that EPS8 enhancement of infection is specific to certain viruses, and the multifunctional nature of EPS8 may impart an anti-viral function for other viruses. In summary, our gene correlation analysis identified both pro- and antiviral host factors with a functional impact on early stages of FLUAV replication without requiring artificial manipulation of the cellular environment. Through interrogation of early steps of FLUAV infection, we established EPS8 as an novel cofactor facilitating FLUAV uncoating.

## ACKNOWLEDGEMENTS

We thank Drs. P Palese, C. Brooke, N. Sherer and Y. Kawaoka for reagents. We thank L. Bollinger for critically editing the manuscript. This work was funded by T32AI078985 to G.P.L., T32GM07215 to V.T., and R01AI125271 and a Shaw Scientist Award to A.M. A.M. holds an Investigators in the Pathogenesis of Infectious Disease Award from the Burroughs Wellcome Fund. This work was also supported in part through Battelle Memorial Institute’s prime contract with the US National Institute of Allergy and Infectious Diseases (NIAID) under Contract No. HHSN272200700016I (Y.C., S.Y., J.H.K.). The views and conclusions contained in this document are those of the authors and should not be interpreted as necessarily representing the official policies, either expressed or implied, of the US Departments of the Army, Defense, and Health and Human Services, or of the institutions and companies affiliated with the authors.

## AUTHOR CONTRIBUTIONS

Conceptualization G.P.L., V.T., S.R.R., J.H.K, A.M.

Methodology, G.P.L., V.T., S.Y., Y.C., C.A.H., D.M.S., S.F.B., S.R.R., J.H.K., A.M.

Formal Analysis, G.P.L., V.T., S.R.R., J.H.K., A.M.

Investigation, G.P.L., V.T., S.Y., Y.C., C.A.H., S.R.R., A.M.

Writing – Original Draft, G.P.L., A.M.

Writing – Review & Editing, G.P.L., V.T., S.Y., Y.C., C.A.H., D.M.S., S.F.B., S.R.R., J.H.K., A.M.

Visualization, G.P.L., V.T., A.M.

Funding Acquisition, G.P.L., V.T., J.H.K., A.M.

Supervision, J.H.K, A.M.

## METHODS

### Cell lines

Authenticated stocks 293T (#CRL-3216), A549 (#CCL-185), and MDCK (#CCL-34) were purchased from the American Type Culture Collection. Parental and edited 293 cells were obtained from Synthego. MDCK-HA cells were a gift from P. Palese (Marsh et al., 2007). All cells were maintained in Dulbecco’s modified Eagle’s medium (DMEM; Corning cellgro) supplemented with 10% heat-inactivated fetal bovine serum (FBS; Atlanta Biosciences) and grown at 37°C with 5% CO_2_. Cells were regularly tested and verified free of mycoplasma contamination using MycoAlert (Lonza, LT07-218). The NCI-60 cell lines are a panel of 59 human breast, central nervous system (CNS), colon, lung, melanoma, ovarian renal cancer, and prostate cancer cell lines (Weinstein, 2006). The NCI-60 panel was obtained from the US National Cancer Institute’s Developmental Therapeutics Program (NCI DTP), Fort Detrick, Frederick, MD, USA. All NCI-60 panel cell lines were grown in RPMI-1640 (ThermoFisher Scientific) supplemented with 10% heat-inactivated FBS (Sigma-Aldrich) at 37°C in a humidified 5% CO_2_ atmosphere. Transfection of 293T and 293 cells was conducted using TransIT-2020 (Mirus, MIR 5400).

### Antibodies

Antibodies used include: anti-EPS8 (BD Biosciences, 610144), anti-M1 (M2-1C6-4R3) (Yewdell et al., 1981), anti-RNP (BEI, NR-3133), anti-tubulin (Sigma, T9026), anti-HSP90 (Santa Cruz Biotechnology, sc-7947), anti-V5 (Bethyl Laboratories, A190-120A), chicken α-mouse AlexaFluor 594 (Invitrogen, A-21201), and donkey α-goat AlexaFluor 488 (Invitrogen, A-11055).

### Viruses

Influenza A virus (FLUAV) strain H1N1 A/WSN/33 (WSN) was propagated in MDCK cells. The recombinant influenza A reporter viruses WSN PASTN (Tran et al., 2013), A/California/04/2009 PASTN (H1N1, CA04 PASTN) (Karlsson, 2015), WSN PASN (Tran et al., 2015), WSN with the polymerase from A/green-winged teal/ OH/175/1983 (H2N1) encoding PB2 S590/R591/K627 (S009-SRK PASTN) (Tran et al., 2015), B/Brisbane/60/2008 (B/Brisbane) PASTN, and FVG-R (Hao et al., 2008) were rescued using the influenza virus reverse genetics system and prepared as previously described. WSN PASN was further purified by centrifugation through a 20% sucrose cushion to remove contaminating luciferase present in the media (Tran et al., 2015). WSN-GFP was amplified and titered on HA-MDCK cells (Marsh et al., 2007).

Multicycle replication infections were performed by inoculating A549 cells at a multiplicity of infection (MOI) of 0.01 using virus diluted in virus growth media (VGM) (DMEM supplemented with penicillin/streptomycin [Corning cellgro], 25 mM HEPES [Corning cellgro], 0.3% BSA [Sigma-Aldrich]) with 0.25 μg/ml TPCK-trypsin. Supernatants were collected at indicated times and titered by plaque assay on MDCK cells (Matrosovich et al., 2006) or by a Nano-Glo viral titer assay by inoculating MDCK cells with reporter viruses and measuring luciferase activity (Karlsson et al., 2018; Tran et al., 2013).

Viral gene expression was measured by infecting cells with WSN PASTN viruses. Virus was diluted in VGM with 0.25-0.5 μg/ml TPCK-trypsin for A549 cells or Opti-MEM I (Invitrogen, 31985070) supplemented with 2% FBS for 293T and 293 cells. Viral gene expression was measured 8 hpi using a Nano-Glo luciferase assay kit (Promega).

Viral attachment was quantified by inoculating A549 cells with bioluminescent PASN virions. Purified virus was diluted in VGM with 0.25 μg/ml TPCK-trypsin, applied to cells for 45 mins at 4°C, and removed. Cells were washed with cold VGM and bound virions were detected by performing a Nano-Glo assay.

FVG-R infections were performed by inoculating A549 cells with virus diluted in Opti-MEM I (Invitrogen, 31985070) supplemented with 0.2% FBS. Viral gene expression was measured 8 hpi using a Renilla luciferase assay system (Promega).

Infections with JUNV (Romero), EBOV, and MARV (Ci67) and infections with RVFV (ZH501) and VEEV (IC-SH3) were conducted under Biosafety Laboratory 4 and 3 conditions, respectively. Cells in 96-well format (30,000 cells per well) were infected at the indicated MOIs. After 1 hour, the inocula were removed, cells were washed with PBS, and replenished with fresh growth media. VEEV and RVFV-infected plates were fixed in formalin 20 hours post-inoculation. All other infected plates were fixed 48 hours post-inoculation. Antigen staining and high-content quantitative image-based analysis were performed as previously described (Radoshitzky et al., 2010, 2016).

### NCI-60 screen and COMPARE analysis

NCI-60 cell lines were seeded by groups of cell origin at 3 × 10^4^ cells per well in 96-well plates (Greiner, 655948 for Operetta; Corning, 3604 for flow cytometry) and grown overnight in RPMI-1640 medium (Lonza, BE12-702F), supplemented with 10% heat-inactivated FBS (Sigma-Aldrich, 4135) at 37°C with 5% CO_2_. The cells were infected with WSN-GFP at MOIs 0.2 and 2. At 3 hours post-inoculation, the cells were washed with RPMI-1640 medium and fresh growth medium was added. WSN-GFP expression was measured by fluorescence microscopy 24 hours post-inoculation. Cells were fixed with 4% paraformaldehyde, and the nuclei were stained with Hoechst 33342 (Thermo Fisher, Catalog: H3570). WSN-GFP virus expression was detected by the Operetta-High Content Imaging System (PerkinElmer Inc.), and the percentage of GFP-positive cells were analyzed by Harmony4.1 software (PerkinElmer Inc.). WSN-GFP expression was evaluated by the flow cytometry (BD Biosciences, LSRFORTESSA) 24 hours post-inoculation. All infections were performed in triplicate, and two biological replicates performed for each MOI condition. Both approaches yielded similar results, and infectivity for each cell line was rank-ordered relative to MDCK cells. The relative infectivity of each cell line was log_2_-transformed and used as input for the COMPARE algorithm (Zaharevitz et al., 2002).

### Knockout and stable expression of EPS8

The *EPS8* locus was edited in A549 cells by lentiviral expression of CRISPR/Cas9 components. Vesicular stomatitis Indiana virus (VSIV) glycoprotein G-pseudotyped lentivirus was generated by transfecting 293T cells with the plasmids psPAX2, pMD2.G, and pLentiCRISPR (Addgene 52961, (Sanjana et al., 2014)) modified to encode a single-guide RNA (sgRNA) targeting *EPS8* (5’-TCAACTTACTTCATCTGAGA-3’, Supplemental Figure 2). A549 cells were transduced with this virus, placed under puromycin selection (0.5 μg/ml), and single cells were cloned. Pooled 293 cells edited at the *EPS8* locus were created by Synthego by transfecting cells with Cas9 RNPs containing an sgRNA targeting exon 5 (5’-GCACTTGACTACCTTTGTCC-3’) (Supplemental Figure 3). 293 cells were single cell cloned. Edited alleles in both cell types were identified by PCR amplification of the locus, Sanger sequencing of the products, and inference of CRISPR edits (ICE) analysis (Hsiau et al., 2019) (Supplemental Figures 2A-B and 3A-B). Knockouts predicted by ICE analysis were assessed by immunoblot. Stable expression of EPS8 in cells was achieved by lentivirus gene delivery. The gene delivery vector pLX304-EPS8 was created by Gateway recombination of pENTR223-EPS8 (DNASU HsCD00505776; (Seiler et al., 2014)) into pLX304 (Addgene 25890) and encodes the 822 amino acid splice variant (NCBI XP_024304650). Virus was produced by transfecting 293T cells with plasmids pLX304-EPS8, psPAX2, and pMD2.G. Wild type and *EPS8*-edited A549 and 293 cells were transduced with this virus and selected with blasticidin to obtain cells stably expressing EPS8.

### Polymerase activity assay

293T cells were transfected with plasmids encoding WSN PA, PB1, PB2, and NP, a vNA-luciferase reporter, a *Renilla* luciferase control reporter, and EPS8 or an empty vector. Firefly luciferase and *Renilla* luciferase activity were assayed 24 hours post-transfection. Firefly luciferase was normalized to *Renilla* luciferase within each sample. Expression of EPS8 was determined by immunoblot of cell lysates.

### Acid bypass assays

Acid bypass with WSN PASTN was performed as described (Matlin et al., 1981; Mondal et al., 2017). Wild type and *EPS8*-edited A549 cells were inoculated at an MOI of 0.1 with virus diluted in VGM with 0.25 μg/ml TPCK-trypsin for 1 hour at 4°C. The inoculum was removed and cells were washed with cold Dulbecco’s phosphate-buffered saline (DPBS). Inoculated cells were then either treated with 20 mM HEPES, pH 7.4 in 154 mM NaCl or 50 mM citrate, pH 5.0 in 154 mM NaCl for 45 seconds at 37°C. The inoculum and treatment buffer were removed and cells were washed with room temperature DPBS. Pre-warmed DMEM supplemented with 10% FBS was added to the cells, infection progressed at 37°C for 8 hours, and viral gene expression was measured by a Nano-Glo assay.

### Immunofluorescence assays

Wild type and *EPS8*-edited A549 cells were grown on coverslips and inoculated with WSN at a MOI of 5 in VGM with 0.25 μg/ml of TPCK-trypsin for 1 hour at 4°C. Warm VGM was added to the cells and infection progressed for the indicated length of time at 37°C. Cells were fixed with 4% paraformaldehyde in DPBS for 30 minutes at room temperature, permeabilized with 0.1% Triton-X in 0.1 M glycine for 5 minutes at room temperature, and blocked in 3% BSA in DPBS for a minimum of 30 minutes at room temperature. Cells were incubated sequentially with primary and secondary antibodies diluted in 3% BSA in DPBS: α-M1 (19 μg/ml) and chicken α-mouse AlexaFluor 594 (2 μg/ml); or α-RNP (1:1000) and donkey α-goat AlexaFluor 488 (2 μg/ml). Coverslips were mounted using mounting medium with 4’,6-diamidino-2-phenylindole (DAPI) stain (Vector Laboratories, H-1200) and imaged using 20X and 40X objectives on an EVOS FL Auto (ThermoFisher). For M1 staining, a minimum of 100 M1-positive cells at 1.5 hpi were counted across 10 random fields of view for each condition in 2 separate biological replicates. Similar quantification was performed at 1 hpi, although fewer M1-positive cells were present for all cell types. RNP localization was quantified by assessing a minimum of 100 cells across 10 random fields of view for each time point in each cell type across 3 separate biological replicates. Images were batch processed using ImageJ for quantification (Schneider et al., 2012). Representative images for cytoplasmic and nuclear RNP staining were batch-processed separately to show staining distribution.

### EPS8 co-immunoprecipitations

Interactions between EPS8 and incoming RNPs was investigated in *EPS8.1* A549 cells stably complemented with EPS8-V5. Cells were inoculated with WSN at an MOI of 25 diluted in cold VGM. Infections were synchronized by inoculating cells at 4°C for 1 hour. Warm VGM was added to the cells and infection progressed for 2.5 hours at 37°C. Cells were washed with cold PBS and lysed in co-IP buffer (50 mM Tris pH 7.4, 150 mM NaCl, 0.5% NP-40) supplemented with protease inhibitors. Lysates were clarified and subjected to immunoprecipitation with 1 μg anti-V5 antibody or control rabbit IgG. Immune complexes were captured with protein A agarose resin, washed extensively with co-IP buffer, eluted, and analyzed by anti-RNP immunoblot to probe for NP.

### Statistics

Each assay was performed in technical triplicate or quadruplicate and represents at least three independent biological replicates with the exception of the immunofluorescence assays which represent at least two biological replicates. Mean and standard deviation were calculated, and statistical significance was tested using a two-tailed Student’s t-test with unequal variance for pairwise comparison or a one-way ANOVA with a Tukey’s honestly significant difference (HSD) *post hoc* analysis for multiple comparisons.

---

**Figure S1.**
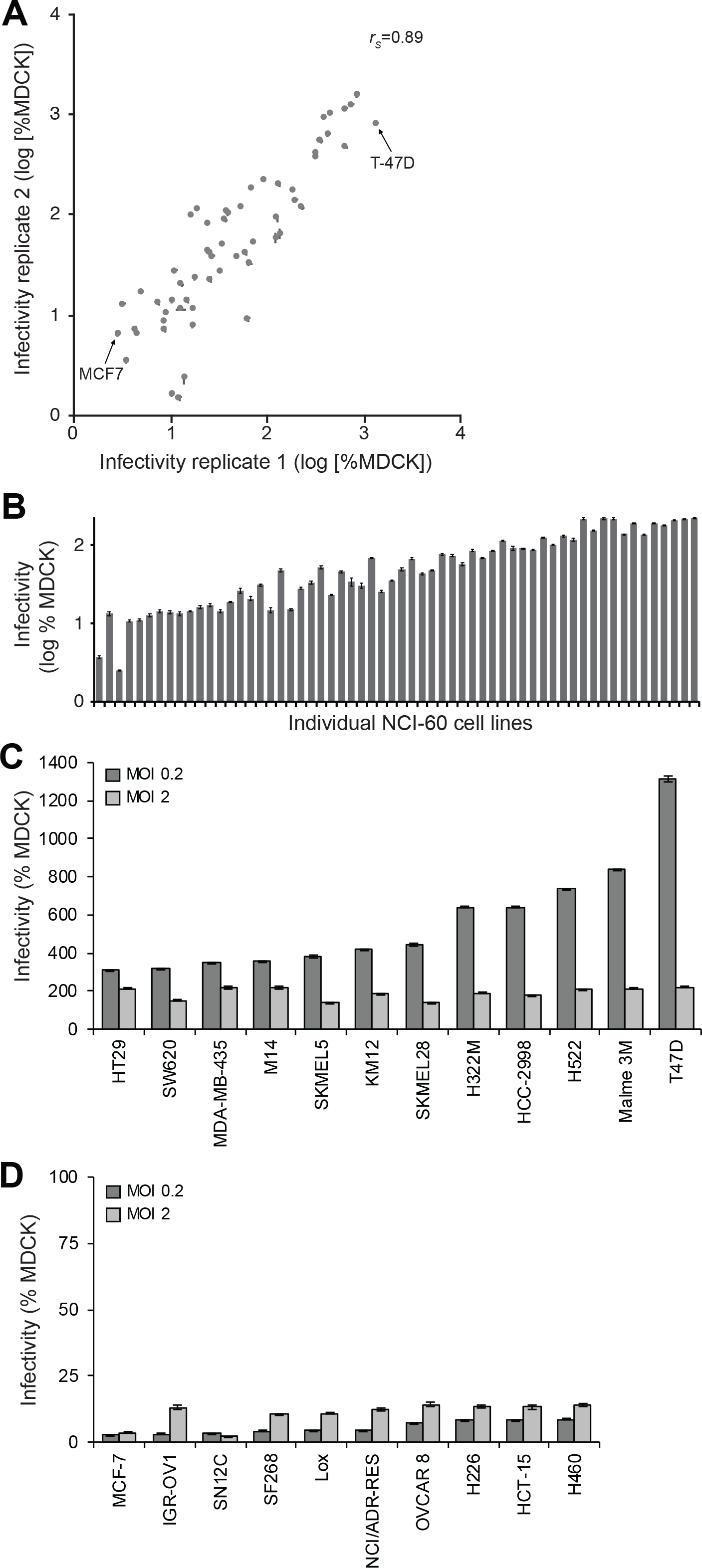
The most susceptible and resistant NCI-60 cell lines to FLUAV infection. The NCI-60 cell lines were challenged with FLUAV in multiple independent screens. (A) Cells were inoculated in replicate screens at an MOI of 0.2 with WSN-GFP and permissiveness to infection was determined by flow cytometry. Data for each cell line were normalized to control MDCK cells, paired across replicates, and plotted against each other. Replicate 1 data are those shown in Figure 1B. (B) Permissiveness to WSN-GFP infection at an MOI of 2 was determined relative to MDCK cells. (C-D) The relative infectivity of the cell lines (C) most resistant to and (D) most susceptible to WSN-GFP infection are shown for screens performed at either a low MOI of 0.2 or a high MOI of 2. For all data, mean of n=3 ± SD, rs= Spearman’s correlation co-efficient.

**Figure S2.**
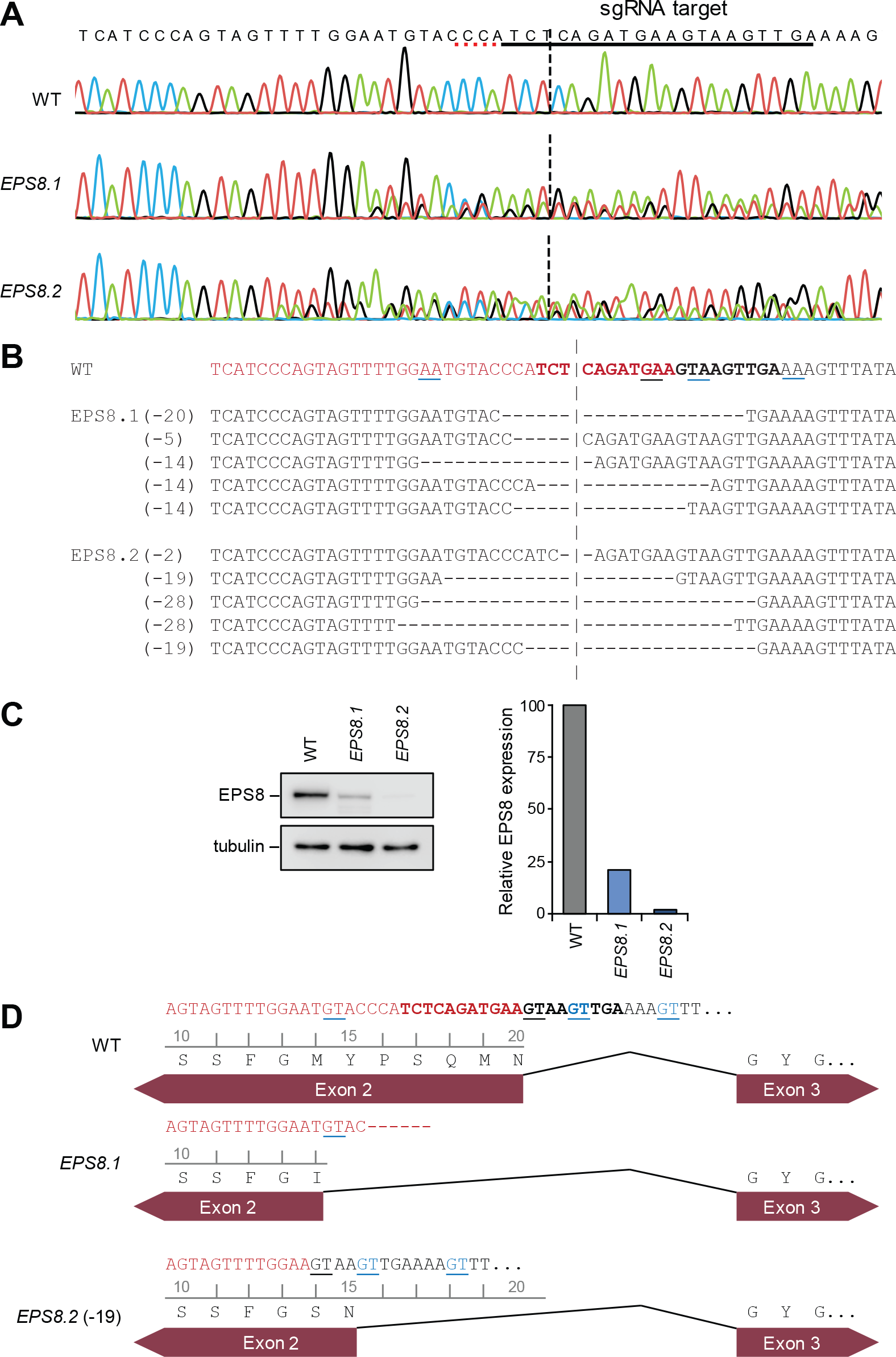
Validation of EPS8 editing in A549 cells. (A) Sequencing chromatograms for wild type and EPS8 knockout A549 cells. The sgRNA target site is underlined in black and the adjacent protospacer adjacent motif (PAM) in red. (B) ICE analysis of edited EPS8 clonal A549 cell lines. All inferred edits contributing at least 5% of the composite genotype are shown. The vertical dashed line indicates the cut site within the sgRNA target (bold letters). The gene exon sequence is shown in red. The two 3’ nucleotides of the native splice donor site are underlined in black and potential alternative splice donors are underlined in blue. (C) Immunoblot of endogenous EPS8 and quantification of band intensity reveal low amounts of EPS8 protein in edited A549 clones despite mutations in the EPS8 locus that should create premature truncations. (D) The indicated deletions could result in low-frequency usage of alternative splice sites to generate a full length EPS8 protein. The wild type gene exon sequence is shown in red. The two 3’ nucleotides of the native splice donor site are underlined in black and potential alternative splice donors are underlined in blue. Hypothetical alternative splice site usage that recreates the EPS8 open reading frame is shown for EPS8.1, where the alternative site matches the consensus splice donor sequence AU/GU, and EPS8.2, where the deletion repositions an alternative site AA/GU that is identical to the native site.

**Figure S3.**
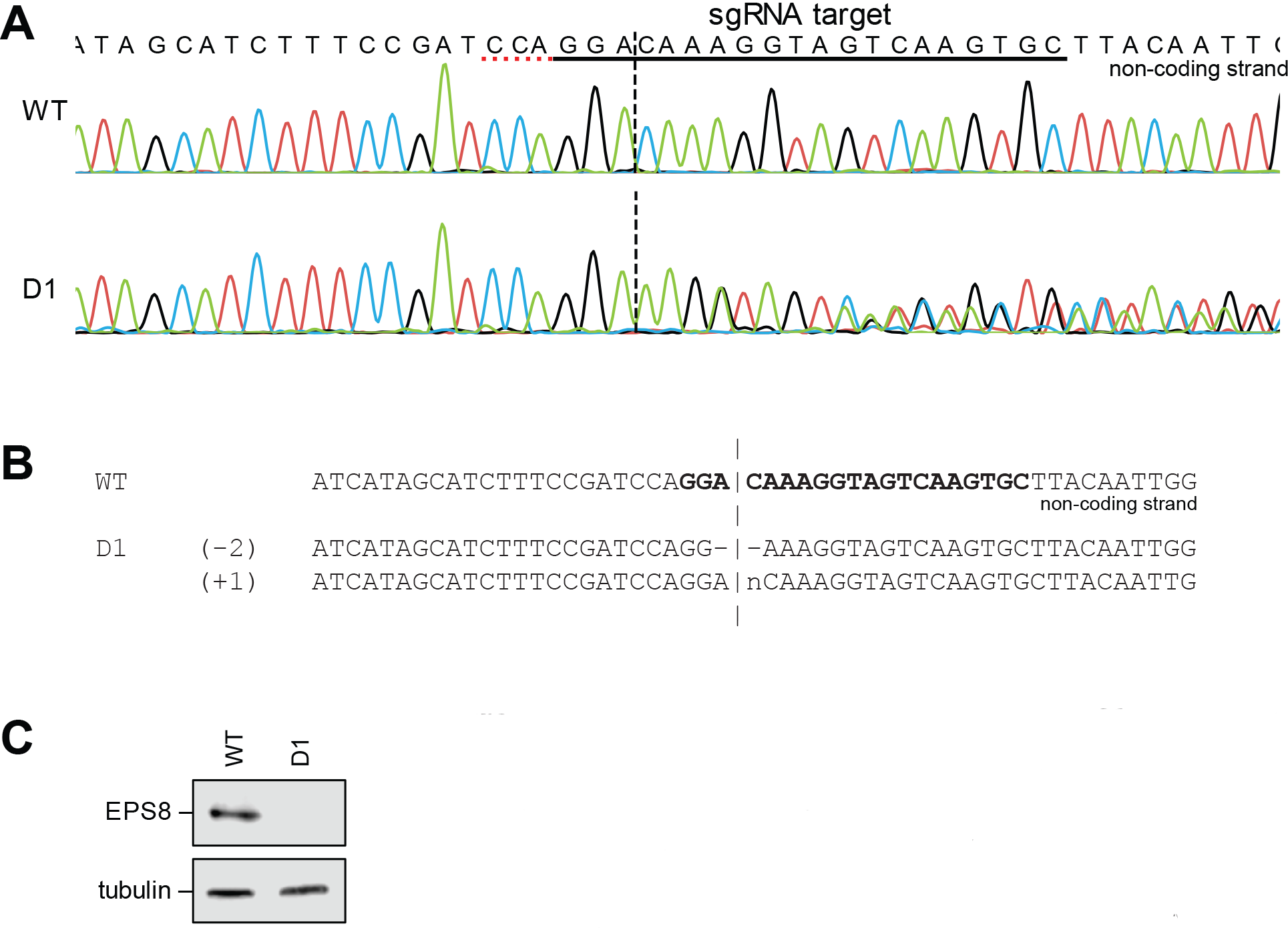
Validation of EPS8 editing in 293 cells. (A) Sequencing chromatograms for wild type and EPS8 knockout 293 cells. The sgRNA target site is underlined in black and the adjacent PAM in red. Note that the non-coding strand is shown. (B) ICE analysis of edited EPS8 clonal 293 cell lines. All inferred edits contributing at least 5% of the composite genotype are shown. The vertical dashed line indicates the cut site within the sgRNA target (bold letters). All edits are predicted to create premature stop codons. (C) Immunoblot of endogenous EPS8 revealed no detectable EPS8 protein in edited 293 cells.

**Figure S4.**
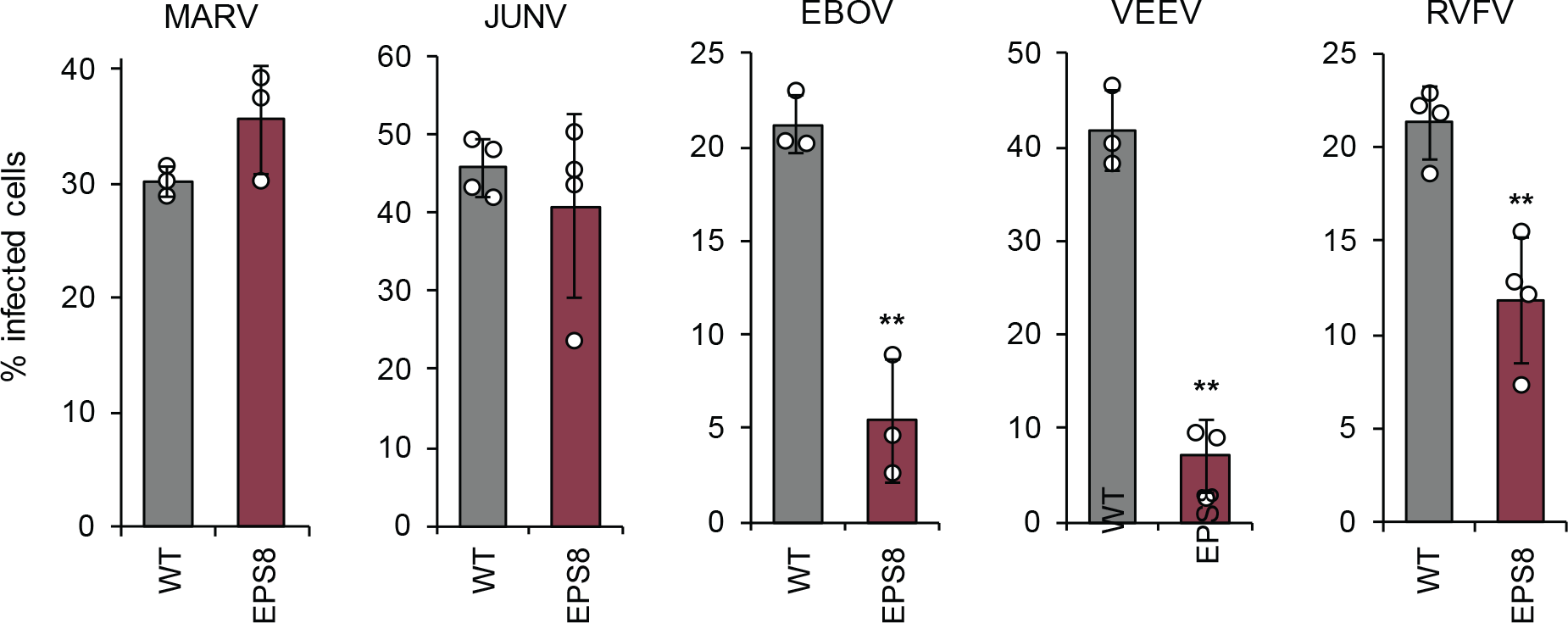
Effects of EPS8 overexpression in A549 on infection by multiple viruses. Wild type A549 cells or A549 cells stably overexpressing EPS8 were infected with Marburg virus (MARV; MOI 5), Junín virus (JUNV; MOI 1), Ebola virus (EBOV; MOI 10), Venezuelan equine encephalitis virus (VEEV; MOI 0.5), or Rift Valley fever virus (RVFV; MOI 1). Cells were stained for the appropriate viral protein and the number of infected (antibody-positive) cells were quantified (mean of n=3-4 ± SD). For all, ** = p<0.01 Student’s two-tailed t-test, unequal variance.

**Figure S5.**
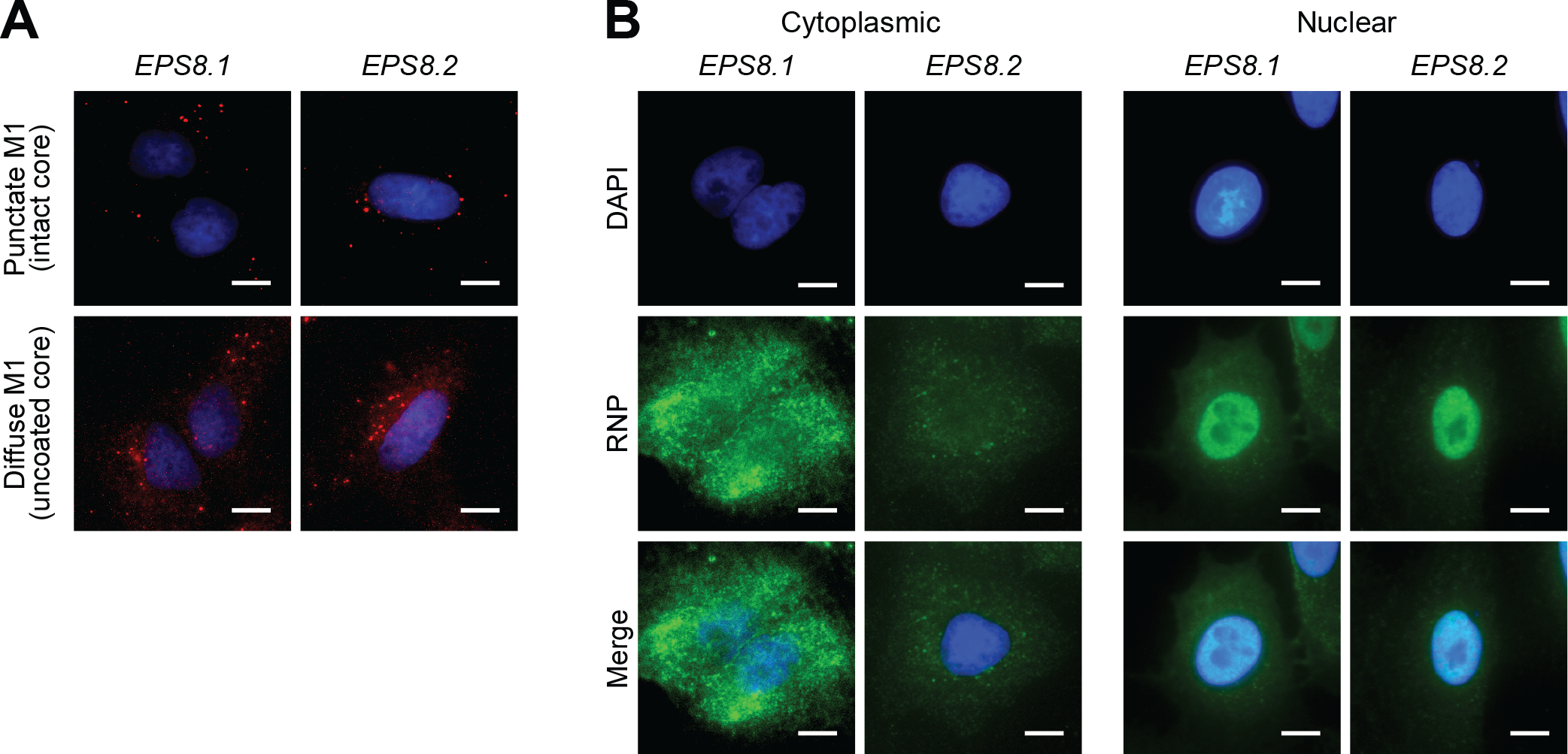
Representative images of cytoplasmic and nuclear RNP staining in EPS8 knockout cells. (A) EPS8-edited A549 cells infected with WSN were stained for M1 (red) and the nucleus (blue). The representative images show punctate and diffuse M1 staining. (B) EPS8-edited cells infected with WSN were stained for viral RNPs (green). Nuclear RNP staining was determined by colocalization of signal with the nucleus (blue). Scale bar = 20 μm.

